# Effect of truffle extracts on growth of *Chlorella vulgaris*

**DOI:** 10.1101/2024.07.03.601813

**Authors:** Anfisa A. Vlasova, Olga E. Lipatova, Alexander Y. Belyshenko, Elena I. Martynova, Natalia A. Imidoeva, Tamara Y. Telnova, Ekaterina V. Malygina, Maria M. Morgunova, Maria E. Dmitrieva, Victoria N. Shelkovnikova, Sophia S. Shashkina, Tatiana N. Vavilina, Denis V. Axenov-Gribanov

## Abstract

Truffles are hypogeous ectomycorrhizal fungi with a complex system of chemical communications and biotic interactions. This study examines the influence of truffle extracts on the growth of the alga *Chlorella vulgaris* as a model organism commonly employed in pharmacology, food industry and biomonitoring. The study involved sampling of fruiting bodies of black truffles in a number of regions across Russia, over a number of years. The observed differences in truffle condition were used to illustrate a potential correlation between the quality and biological activity of truffles. Methanolic extracts were prepared and added to the culture of *C. vulgaris* to evaluate the growth of algae using the spectrophotometry methods. For the first time, we demonstrated that truffle extracts had biological activity in terms of the algae *C. vulgaris* growth stimulation. We observed the effects of short-term and long-term growth stimulation. To date, it can be concluded that there is no direct correlation between the stimulating effects on algae and the state of truffles, place of their sampling, or their quality. Finally, it is crucial to examine biological regulations that operate in complex systems such as truffles. It is suggested that in the near future truffles may become a model system for the study of complex biotic and chemical communications.

## 1. INTRODUCTION

Truffles are a worldwide famous gastronomy delicacy. Truffles are known for their flavor, cosmetic, and aphrodisiac properties. They are also known as a source of biologically active compounds and natural products that exhibit antiviral, antimicrobial, antimutagenic, antioxidant, and anti-inflammatory activities [1]. One of the most notable characteristics of truffles is their flavor. Each species of truffle produces a unique blend of flavors that includes alcohols, ketones, aldehydes, aromatic compounds, and other compounds. Differences in flavor profiles of different truffle species arise from both abiotic and biotic factors. Biotic factors include bacteria, fungi, host plants, etc. [2]. Abiotic factors are climate, weather and other environmental parameters.

Being the hypogenic ectomycorrhizal fungi, truffles participate in symbiotic relationships with microorganisms, plants, and animals. Truffle spores are spread by mycophagous animals, who consume the fruiting bodies of truffles and ferment them in their gastrointestinal tract [3,4,5]. The effects and impacts of truffles on mycophagous animals have not been studied. Once released from the gastrointestinal tract of animals, spores develop into mycelium, which forms the ectomycorrhizal association with the roots of host trees [5,6,7,8]. Next partners in the symbiosis between the fungus and the host tree are bacterial communities [9,10]. Fruiting bodies of truffles are a home for many bacterial species, whose role in the fungal association is unclear. For example, bacteria *Bradyrhizobium* sp. are the main truffle-associated bacteria involved with the truffle species studied [10,11,12]. Bacteria associated with fruiting bodies of truffles often act as assistants of mycelial growth, adaptation, forming of fruiting bodies, and other processes [10]. Also, symbiont bacteria can synthesize volatile organic compounds that contribute to the truffle flavor [13,14]. The VOCs synthesized by truffle symbiont bacteria also play an important role in the interactions with other organisms [15].

In addition to the association between truffles and bacteria, cyanobacteria and microalgae are also regular members of the truffle consortium [16,17]. One of well-studied groups of microalgae belongs to genus *Chlorella*.

*Chlorella* is widely used as a source of pharmacologically active metabolites with different biological activities [18,19]. This alga is also being studied for couplings with other organisms, namely bacteria and viruses. For example, it has been shown that *Azospirillus* sp. and *Bacillus* sp. can stimulate growth of *C. vulgaris*. In the study by Cho K. (2019), it was shown that co-cultivation of algae and bacteria stimulates the growth of *C. vulgaris* and increases its biomass [20].

The aim of our study was to determine the biological activity of black truffles collected from different regions against the green alga *Chlorella vulgaris*.

## 2. MATERIALS AND METHODS

### 2.1. Sampling of truffles

Wild black truffles were sampled in several cities of the Southern region of the Russian Federation between August and November in 2020□2022. The samples were taken near the cities of Krasnodar, Maykop, and Sochi.

Fungi sized 3 to 5 cm were collected and cleaned with a soft toothbrush and then sterilized with 70% ethyl alcohol. Then, the truffles were dried and shock-frozen in liquid nitrogen. Until further experiments, the samples were stored in liquid nitrogen.

### 2.2. Extraction

To perform the extraction, 0.5 grams of each truffle sample were defrosted and homogenized using a mortar and pestle with methanol added in a ratio of 1:10. The mixture was incubated and shaken for one hour on a roller shaker MX-T6-S (China). Then, the samples were centrifuged at 3 000 rpm for 10 minutes using DM 0412 centrifuge (Hettich, USA). The extracts were dried *in-vacuo* and re-dissolved in methanol in the concentration of 10 mg/mL.

Each truffle extract was assigned a unique identification number (Table 1).

**Table 1.** List of extracts, obtained from mushrooms and tested in this study.

| Place of collection | Extract ID | Quality of truffles |
| --- | --- | --- |
| Krasnodar 2020 | *Tuber* sp. LPB2020-16 | Fresh, firm, faint odor |
|  | *Tuber* sp. LPB2020-17 | Fresh, firm, faint odor |
| Krasnodar 2021 | *Tuber* sp. LPB2021-26 | Fresh, firm, specific odor |
|  | *Tuber* sp. LPB2021-27 | Fresh, firm, specific odor |
|  | *Tuber* sp. LPB2021-28 | Fresh, firm, specific odor |
|  | *Tuber* sp. LPB2021-29 | Fresh, firm, specific odor |
|  | *Tuber* sp. LPB2021-30 | Fresh, firm, specific odor |
| Krasnodar 2022 | *Tuber* sp. LPB2022-67 | Stale, soft, specific odor |
|  | *Tuber* sp. LPB2022-69 | Stale, soft, specific odor |
|  | *Tuber* sp. LPB2022-39 | Fresh, firm, specific odor |
|  | *Tuber* sp. LPB2022-42 | Fresh, firm, specific odor |
|  | *Tuber* sp. LPB2022-47 | Stale, soft, sharp unpleasant odor |
| Sochi 2021 | *Tuber* sp. LPB2021-22 | Fresh, firm, specific odor |
|  | *Tuber* sp. LPB2021-23 | Fresh, firm, specific odor |
|  | *Tuber* sp. LPB2021-24 | Fresh, firm, specific odor |
| Maykop 2022 | *Tuber* sp. LPB2022-58 | Fresh, firm, specific odor |
|  | *Tuber* sp. LPB2022-59 | Stale, soft, sharp unpleasant odor |
|  | *Tuber* sp. LPB2022-61 | Fresh, firm, specific odor |

### 2.3. Assessment of truffle extract effects on growth of algae *C. vulgari*s

In this study, the biological activity of extracts was evaluated on the basis of growth of *C. vulgaris*. First, the mother culture of algae was grown in 50% of Tamiya nutrient medium at 36 □ for 5 days. Then, the culture was diluted in 50% of Tamiya nutrient media until the optical density value reached 0.06-0.08 at the wavelength of 560 nm. General technique of spectrophotometric analysis was applied using PE-5300V spectrophotometer (PromEcoLab, Russia). Composition of 100% Tamiya nutrient medium is as follows: KNO_3_ - 5.0 g/L, MgSO_4_ x 7H_2_O - 2.5 g/L, KH_2_PO_4_ x 3H_2_O - 1.25 g/L, FeSO_4_ x 7H_2_O - 0.009 g/L, H_3_BO_3_ - 2.86 g/L, MnCl_2_ x 4H_2_O - 1.81 g/L, ZnSO_4_ x 7H_2_O - 0.22 g/L, MoO_3_ x 4H_2_O – 0.018 g/L, NH_4_VO_3_ - 0.023 g/L.

For the experiment, aliquots of methanol extracts in amount of 100 µl were added to conical flasks. The solvent was evaporated, and then, 40 mL of the *C. vulgaris* culture was added to the dry residue.

The control group of *C. vulgaris* was cultivated with dry residue of methanol added, as a solvent of truffle extract. Truffle extracts were not added to the samples.

The experiment was conducted in three stages. In the first stage, the experimental group included the following extracts: *Tuber* sp. LPB2020-16, *Tuber* sp. LPB2020-17, *Tuber* sp. LPB2021-22, *Tuber* sp. LPB2021-23, *Tuber* sp. LPB2021-24, *Tuber* sp. LPB2021-26, *Tuber* sp. LPB2021-28, *Tuber* sp. LPB2021-29, and *Tuber* sp. LPB2021-27. In the second stage, the experimental group included the following extracts: *Tuber* sp. LPB2022-58, *Tuber* sp. LPB2022-59, *Tuber* sp. LPB2022-61, *Tuber* sp. LPB2022-67, and *Tuber* sp. LPB2022-69. In the third stage, the experimental group included the following extracts: *Tuber* sp. LPB2021-30, *Tuber* sp. LPB2022-39, *Tuber* sp. LPB2022-42, and *Tuber* sp. LPB2022-47. Each stage had its own control group of *C. vulgaris*.

Further, control and experimental groups were cultivated at 36 □using whirlpool photothermoshaker x12-1 (Mycotech, Russia). The cultivation time was up to 6 days with 900 Lux light flow. To assess the dynamics of the algae growth, we measured the culture optical density on days 2, 4 and 6 of the experiment.

### 2.4. Statistical Analysis

In total, we estimated the effects of 18 fruiting bodies of truffles. All experiments were carried out in 3–7 biological replicates. Statistical analysis was performed with Past software (V. 4.03) using Tukey’s Proc ANOVA test. Tukey’s range test, p-value of ≤0.05 was considered to indicate statistical significance.

## 3. RESULTS

During the experiment, growth of *C. vulgaris* was estimated on days 2, 4 and 6 of the experiment.

Figure 1 demonstrates the growth of *C. vulgaris* when methanol extracts of truffles were added. Also, Table 1 shows the dynamics of growth of *C. vulgaris* algae (expressed in % related to control group).

**Fig.1.**
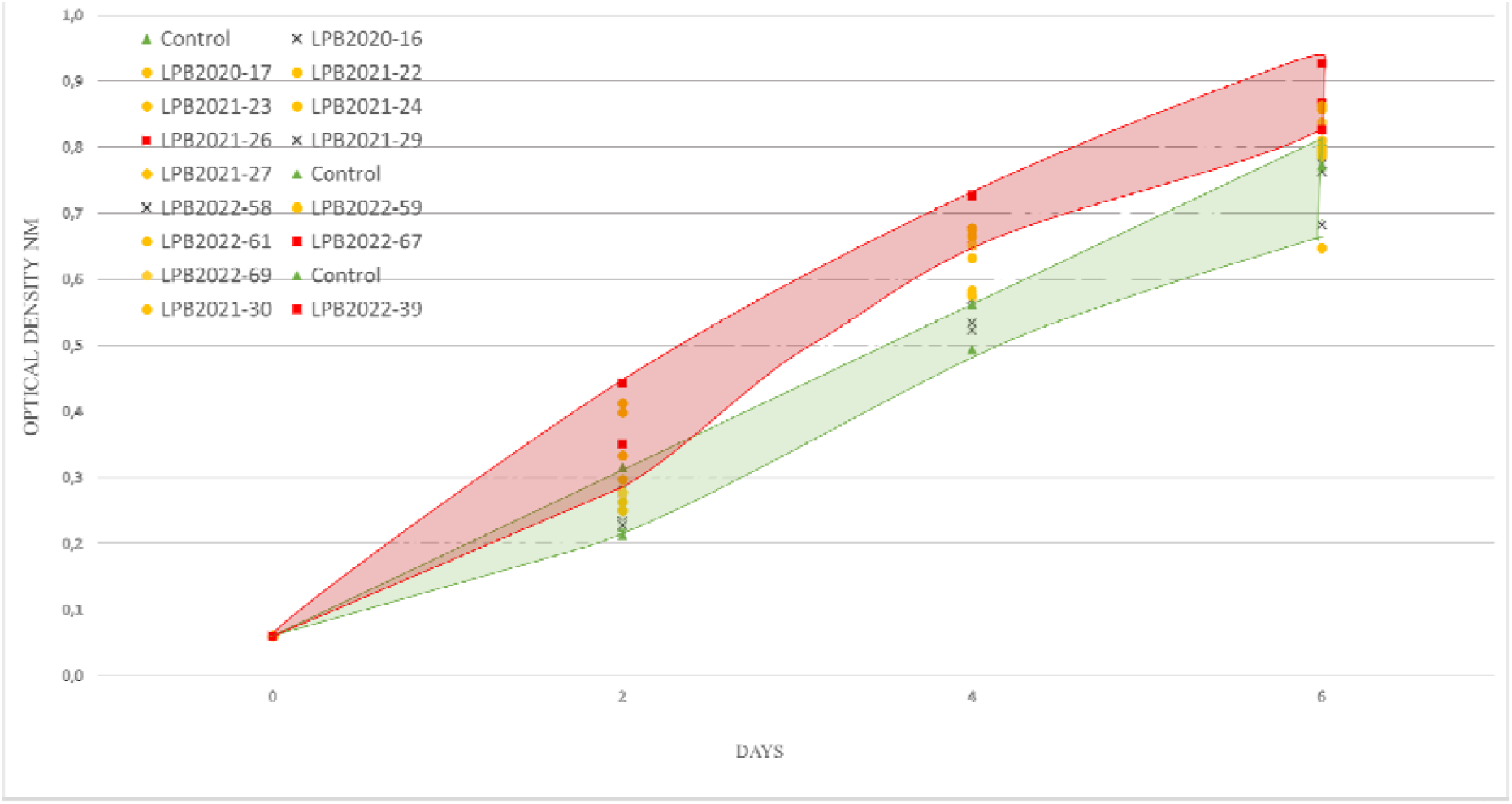
Optical density of *C. vulgaris* under control and experimental conditions. Legend: 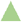- Control; 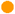 - significant differences observed in one or two control points (short- term effects, i.e. after 2, 4, or 6 days of the experiment); 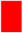-significant differences observed during 6 days of experiment (long-term effects); ⍰- no significant differences. P-value <0.05

Green area in Figure 1 indicates the area of the *C. vulgaris* control group cultivated without adding truffle extracts. Red area shows the zone where the experimental group showed notable differences from the control one, throughout the experiment (long-terms effects). Yellow markers show the short-term stimulation effects on the algae, but significant differences from the control group can be seen in one or two control points. Gray markers indicate extracts that did not stimulate growth of *C. vulgaris*.

Four out of 18 truffle extracts exhibited the highest long-term stimulating effect on algal growth. During 6 days of the experiment, we observed significant stimulating effects of the following methanol extracts: *Tuber* sp. LPB2021-26 (sampling: Krasnodar 2021), *Tuber* sp. LPB2022-67 (sampling: Krasnodar 2022), *Tuber* sp. LPB2022-39 (collection: Krasnodar 2022), and *Tuber* sp. LPB2022-47 (sampling: Krasnodar 2022). After adding *Tuber* sp. LPB2021-26 extract to the culture of *C. vulgaris*, we observed the increase of the algal culture optical density. Optical density of the algal suspension increased from 0.06 to 0.352 on the second day of the experiment, 0.494 on the fourth day, and 0.868 on the sixth day. Compared to control conditions, in the experimental group, optical density of the algal culture was 56%, 36% and 26% higher on the 2nd, 4th and 6th day of the experiment, relatively. In the case of the effect of *Tuber* sp. LPB2022-67 extract, the daily growth reached 30% in the first two days of experiment and then decreased to the end of experiment (Table 1).

Also, it was found that some extracts exerted partial stimulation (short-term stimulation effects) of algal growth. Significant differences between control and experimental groups were observed at one or two points in time. For example, truffle extract *Tuber* sp. LPB2021-22 demonstrated significant differences on days 4 and 6 of the experiment. Growth of *C. vulgaris* comprised 7.5% to 8.5% per day. Extracts of *Tuber* sp. LPB2021-23 and *Tuber* sp. LPB2021-24 stimulated the algae growth and demonstrated significant differences compared to control conditions during the first four days of the experiment. On the second day, the optical density of *Tuber* sp. LPB2021-24 extract was 0.298, which was 32% higher than under control conditions. On day 4, optical density was 0.678 (37% rate of stimulation).

The *Tuber* sp. LPB2021-23 extract revealed the highest short-term stimulation activity towards the algae. Optical density of *Tuber* sp. LPB2021-23 extract at the 2-day point was 0.399, which was 77% higher related to the control conditions. However, by day 6 of the experiment, the effects of the truffle extract decreased, and optical density of algal culture was similar to the control value.

The study revealed that adding extracts of *Tuber* sp. LPB2020-16 (sampling: Krasnodar 2020), *Tuber* sp. LPB2021-29 (sampling: Krasnodar 2021), and *Tuber* sp. LPB2022-58 (sampling: Maykop 2022) did not stimulate algal growth.

## 4. DISCUSSION

In our study, we demonstrated for the first time that the extracts obtained from black truffles impose growth stimulation effects on microalgae *C. vulgaris*. For this study, the fruiting bodies of truffles were sampled in a number of regions across Russia, over a number of years. The observed differences in the state of truffles were used to reveal a potential correlation between the quality and biological activity of truffles. To date, we can conclude that there is no direct correlation between the stimulating effects on algae and the state of truffles, place of their sampling, or their quality. At the same time, several reactions of C. vulgaris were observed when cultivating it with truffle extracts. Here, we detected both long-term and short-term stimulation effects that truffle mushroom extracts exert on the algae growth.

We demonstrated that the highest stimulating effects of extracts occurred during the first two days of the experiment. Subsequently, all types of truffle extracts retained their stimulating effect on *C. vulgaris*, but a decrease in growth rate was noted (Table 2). This could be explained by composition and energetic value of the nutrient media that decreased during the experiment. Another explanation could be that over time *C. vulgaris* caused a thinning of the culture medium and reduced the amount of stimulating natural products in the truffle extracts. In this case, the variable effects of our extracts may be related to unstable chemical composition of the truffles due to unstable content of microflora associated with them.

**Table 2.** Daily growth dynamics of the alga *C. vulgaris*.

| Growth dynamics of the alga *C. Vulgaris* |  |  |  |
| --- | --- | --- | --- |
|  | 2-day | 4-day | 6-day |
| LPB2020-16 | 2 | 3 | 7 |
| LPB2020-17 | 8 | 9 (*) | 9 (*) |
| LPB2021-26 | 28 (*) | 18 (*) | 13 (*) |
| LPB2021-27 | 24 (*) | 7 (*) | -3 |
| LPB2021-28 | 26 (*) | 13 (*) | 8 |
| LPB2021-29 | 12 | 4 | 0 |
| LPB2021-30 | 15 (*) | 10 (*) | 2 |
| LPB2022-67 | 15 (*) | 7 (*) | 7 (*) |
| LPB2022-69 | 15 (*) | 7 (*) | 2 |
| LPB2022-39 | 20 (*) | 15 (*) | 4 (*) |
| LPB2022-42 | 6 (*) | 3 | 2 |
| LPB2022-47 | 5 (*) | 8 (*) | 6 (*) |
| LPB2021-22 | 6 | 9 (*) | 8 (*) |
| LPB2021-23 | 39 (*) | 14 (*) | 8 |
| LPB2021-24 | 16 (*) | 19 (*) | 13 |
| LPB2022-58 | 4 | 0 | -3 |
| LPB2022-59 | 9 (*) | 0 | 0 |
| LPB2022-61 | 15 (*) | 8 (*) | 3 |

Legend: asterisks indicate significant values. Pink color means long-term stimulation effects; Yellow color — short-term effects. White color — no effects

In this study, we observed that extracts that stimulated algal growth were obtained from truffle fruiting bodies and collected at different time. No patterns of change in algal growth were detected in this case. Thus, *Tuber* sp. LPB2021-26 extract was isolated from truffles collected in 2021, while extracts of *Tuber* sp. LPB2022-67, *Tuber* sp. LPB202239, and LPB2022-47 were isolated from fruiting bodies collected in 2022.

The same conclusion has been made for extracts that exhibited short-term stimulation effects. They were also collected in a number of years in different regions of Russia. For example, *Tuber* sp. LPB2021-30 extract was obtained from a fruiting body collected in 2021 near Krasnodar. Extract of *Tuber* sp. LPB2022-61 was obtained from a truffle sampled in 2022 near Sochi. The dynamics of the algae growth was similar when adding these extracts.

It is important to note that the extracts of fruiting bodies imposed the same effect on growth of *C. vulgaris* regardless of the quality of truffles. For example, the extract obtained from *Tuber* sp. LPB2021-26 (fresh and firm truffle with a characteristic odor) had the same long-term stimulation effect as the extract from *Tuber* sp. LPB2022-47 (stable and unstable truffles with sharp, unpleasant odor).

Thus, the mechanisms of influence of truffles on algae or plants are understudied. To date, we found only one study related to regulation of plant root morphogenesis by truffles. This study showed that at early stage of fungal-plant interaction, *Tuber borchii* and *Tuber melanosporum* induced changes in the root morphology of *Cistus incanus* and *Arabidopsis thaliana* due to the production of indole-3-acetic acid and ethylene by the mycelium [21]. This paper only described the influence of truffle mycelium on plants, and did not address the effects of a complex mixture of natural products produced by truffle fruiting bodies and symbiotic microorganisms.

As mentioned above, activity of the extracts can be related to the microflora inhabiting truffles. The life cycle of truffles includes symbiosis with not only host plants, but also with microorganisms. Symbiont bacteria exert their influence not only on truffles themselves, but also on the surrounding organisms, too. Therefore, we hypothesize that it was specific symbiont bacteria that had the resulting effect on the growth of the alga *C. vulgaris*. For example, a strain of *Staphylococcus aureus* has been shown to synthesize volatile organic compounds potentially involved in inhibiting the growth of *T. borchii* mycelium [12,22]. It has been reported that in pure culture, the isolates of *Pseudomonas* spp. obtained from *T. borchii* ascoma are capable of producing phytoregulatory and biocontrol substances that influence the growth and morphogenesis of *T. borchii* mycelium [12,23]. Volatile organic compounds produced by microbes can influence plant adaptability, they can also interfere with plant-animal interactions. By doing so, these compounds promote plant growth and prevent the emergence of harmful microbes [12,22,24]. It has been shown that *Azospirillum* sp. and *Bacillus* sp. can stimulate growth of *C. vulgaris* [25,20]. It was found that selective inoculation of nutrient medium with bacteria promoted growth of the alga *C. vulgaris* and increased its biomass [20]. This may suggest that the volatile organic compounds produced by truffle symbiont bacteria may stimulate growth of *C. vulgaris*. The microbial composition of Russian black truffles collected in Sochi has been described earlier in our studies [10]. The primary analysis detected at least 45 bacterial organisms with unique OTUs, and 22 fungal organisms as regular inhabitants of *Tuber* sp. fruiting bodies [10,9,13,26].

Different composition of symbionts of truffles, influencing the content of natural products in the metabolome of truffles, allows us to hypothesize that both truffles and microalgae could have common natural products in their metabolomic pathways. These natural products can be found in literature, they are 1-octen-3-ol (eight-carbon-containing volatiles) and 2-methyl-1-propanol. As predicted before, these natural products have been detected in most truffles species and might act as signals to plants [27]. Also, 1-octen-3-ol previously was found in *Chlorella* sp., as a product of linoleic acid oxidation and was described as a natural product responsible for aroma of seafood [28,29,30].

Thus, this study is one of the first to assess the stimulating effects of true truffles. Microbial associations contribute significantly to the biological activity of truffles and cause non-stable effects. However, complicated symbiotic couplings between a truffle and other organisms, such plants and microorganisms, require the study of biological regulation in these complex systems. Truffles may become a model system for the investigation of complex biotic and chemical communications in the near future.

The study was conducted with the support of the Russian Science Foundation (project 22-76-10036).

